# Intraoperative arteriovenous patient sampling to assess *in situ* non-small cell lung cancer metabolism

**DOI:** 10.1101/2025.08.05.668488

**Authors:** Johnathan R. Kent, Keene L. Abbott, Rachel Nordgren, Amy Deik, Nupur K. Das, Millenia Waite, Tenzin Kunchok, Anna Shevzov-Zebrun, Nathaniel Christiansen, Amir Sadek, Darren S. Bryan, Mark K. Ferguson, Jessica S. Donington, Alexander Muir, Yatrik M. Shah, Clary B. Clish, Matthew G. Vander Heiden, Maria Lucia L. Madariaga, Peggy P. Hsu

## Abstract

We performed intraoperative arteriovenous sampling in participants undergoing surgical resection for non-small cell lung cancer (NSCLC) to characterize in situ tumor metabolism, directly measuring metabolite consumption and secretion in tumor-bearing lung versus normal lung and the systemic circulation. Healthy lung tissue secreted lower levels of lactate, pyruvate, and tricarboxylic acid (TCA) intermediates and consumed less glucose compared to the systemic circulation. In contrast, tumor-bearing lung demonstrated elevated lactate secretion, along with increased efflux of succinate, fumarate, glycine, and aspartate, despite similar glucose uptake. Lactate secretion correlated with tumor PET avidity but not size, and overall metabolic profiles distinguished cancerous from normal lung tissue. These findings confirm enhanced glycolysis in NSCLC in vivo, while also revealing context-dependent patterns of TCA metabolite accumulation and amino acid secretion. Our results demonstrate the utility of intraoperative sampling to uncover metabolic features of human tumors.

## Main Text

Alterations in metabolism are a hallmark of cancer (1). Carl and Gerty Cori, contemporaries of Otto Warburg, performed seminal experiments a century ago on chickens with tumors implanted unilaterally in one wing (2). By sampling the veins draining each wing, they compared the metabolic profiles of tumor-bearing and non-tumor-bearing tissue. They found lower glucose levels and increased lactate in the blood from the vein draining the tumor-bearing wing, data supporting Warburg’s description of aerobic glycolysis in cancer.

Recent studies in cancer metabolism have employed stable isotope tracing and measurement of metabolite levels in tissues, both techniques built upon improvements in using mass spectrometry to assess metabolism. While powerful, these approaches are limited in their ability to directly measure metabolite consumption and secretion. Furthermore, human studies often rely on cancer cell lines, whose metabolism can be influenced by media conditions, or on surgical specimens, where metabolic integrity is compromised by prolonged arterial ligation. We sought to gain deeper insights into lung cancer metabolism by intraoperative sampling, which avoids these pitfalls, in a modernized Cori experiment.

We directly evaluated non-small cell lung cancer (NSCLC) metabolism by sampling the arterial inflow and venous outflow of tumor-bearing versus non-tumor-bearing lung tissue in patients undergoing lung cancer resection (**Supplemental Figure 1A-E**). This intraoperative arteriovenous sampling enabled determination of how metabolite concentrations change in venous compared to arterial circulation involving the diseased and non-diseased lung, as well as in the patient-matched systemic circulation. Blood samples were obtained during 20 surgical resections (**Supplemental Table 1**), with a mean tumor size of 3.13 cm. Most tumors were adenocarcinomas (90%) and half represented stage 1 disease (**Figure 1A**); 25% of participants had received neoadjuvant therapy (Chemotherapy: 1; Chemotherapy + Immunotherapy: 2; On clinical trial with chemotherapy +/- Immunotherapy: 2). At the time of surgery, after dissection of the hilar vessels, blood samples were obtained from the pulmonary artery (arterial inflow) and pulmonary veins (venous outflow; tumor-bearing and normal lobes). Simultaneously blood samples were obtained from a radial arterial line and a peripheral vein to provide an approximation of the concentration gradient change of metabolites through the systemic circulation (**Supplemental Figure 1A-E**). All samples were kept cold and processed within minutes. To assess sample stability, we separately mimicked intraoperative processing time and handling conditions and found that >97% of polar metabolites remained unchanged during processing (**Supplemental Figure 1F-H**). No intraoperative bleeding or complications from pulmonary vasculature sampling occurred. Seven participants suffered a postoperative complication (4 – atrial fibrillation, 2 – prolonged air leak, 1 – acute respiratory distress syndrome & mortality) unrelated to this study.

**Figure 1.**
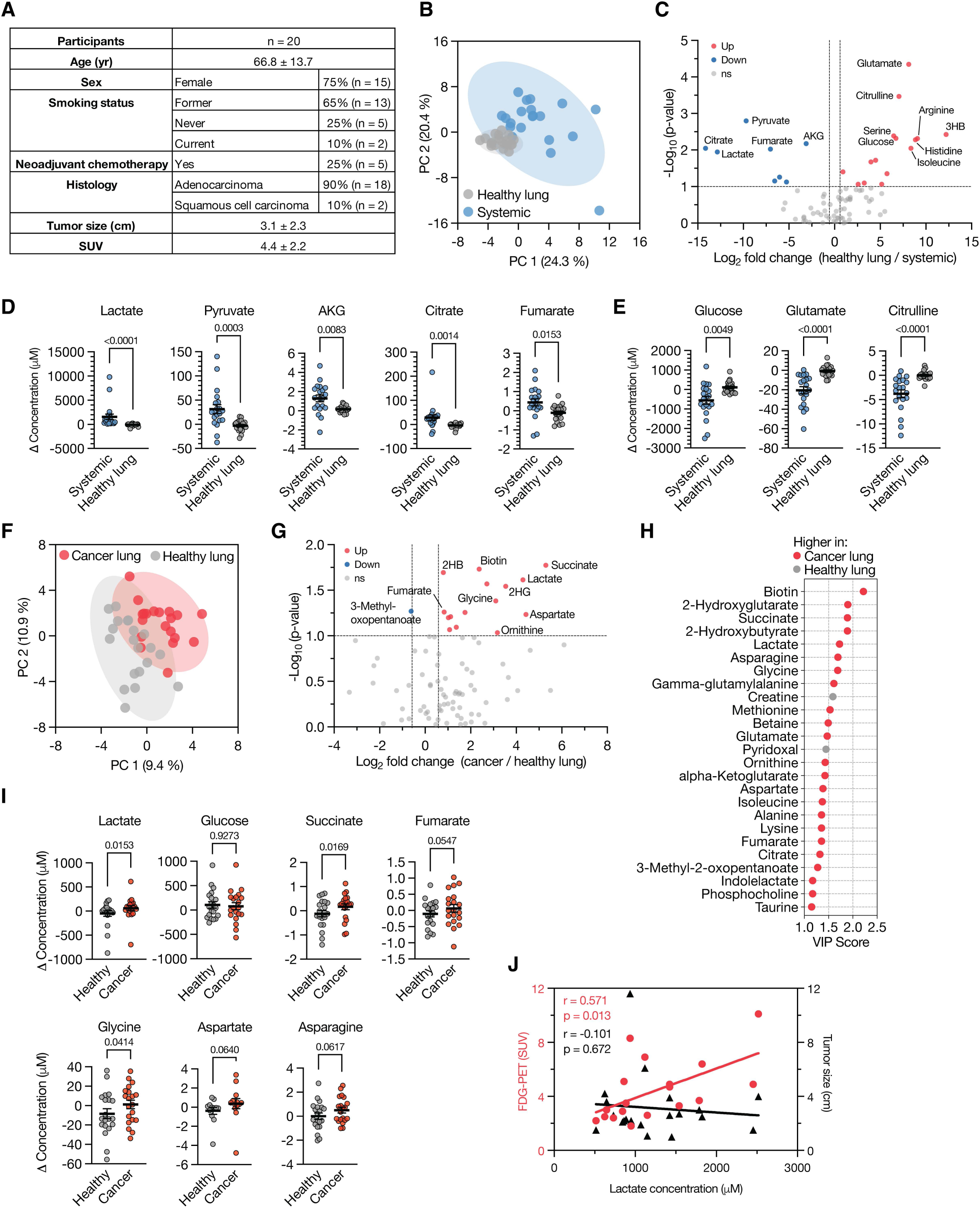
Intraoperative arteriovenous sampling reveals nutrient consumption and production in healthy versus tumor-bearing lung tissue. **(A)** Summary of clinical characteristics for the 20 NSCLC participants included in the study. **(B)** Partial least squares discriminant analysis (PLS-DA) comparing metabolite profiles of blood derived from analysis of non-tumor-bearing (healthy) lung circulation (pulmonary vein of a non-tumor lobe minus pulmonary artery) versus systemic circulation (peripheral vein minus radial artery). **(C)** Volcano plot showing metabolites significantly different when comparing metabolite changes across the healthy lung and the systemic circulation. Significance was defined as a fold change >1.5 and a raw p-value <0.1 (paired two-tailed t-test, n = 20). **(D-E)** Selected metabolites significantly elevated (D) or reduced (E) in systemic circulation compared to healthy lung. Data are presented as mean ± SEM. P-values calculated using the Wilcoxon matched-pairs signed rank test (n = 20). **(F)** PLS-DA comparing blood metabolite profiles from analysis of cancer-bearing lung circulation versus healthy lung circulation (both calculated as pulmonary vein minus pulmonary artery). **(G)** Volcano plot depicting metabolites significantly different when comparing metabolite changes across the cancer-bearing and the healthy lung. Thresholds and statistics as in panel (C); n = 20. **(H)** Variable importance in projection (VIP) plot of the top 25 metabolites distinguishing cancer and healthy lung. Metabolites in red are elevated in cancer lung; those in grey are elevated in healthy lung. Derived from data shown in (F). **(I)** Selected metabolites differing in circulation across cancer-bearing and healthy lung. Data are mean ± SEM; p-values from Wilcoxon matched-pairs signed rank test (n = 20). **(J)** Scatter plot showing lactate levels in the cancer-draining pulmonary vein plotted against FDG-PET SUV (red, n = 18) or tumor size (black, n = 20). Pearson correlation coefficients and p-values are indicated. 2HB: 2-hydroxybutyrate; 3HB: 3-hydroxybutyrate; AKG: alpha-ketoglutarate.

In each sample collected we performed quantitative polar metabolomics using liquid chromatography/mass spectrometry (LC/MS) with a library of chemical standards as previously described (3). To establish a baseline for normal lung metabolism, we compared metabolite concentration changes in the systemic circulation (peripheral vein vs. radial artery) to those in the non-tumor-bearing (healthy) lung (pulmonary vein of a non-tumor lobe vs. pulmonary artery). Principal component analysis revealed separation in metabolite changes between systemic and lung-derived blood (**Figure 1B**), with many metabolites showing differential utilization (**Figure 1C, Supplemental Figure 2A**). Specifically, healthy lung tissue secreted lower levels of metabolites such as lactate, pyruvate, α-ketoglutarate, citrate, and fumarate, and consumed less glucose, glutamate, and citrulline compared to that observed in the systemic circulation (**Figure 1D-E**). These findings align with prior studies suggesting that, compared to other organs, the lung is relatively metabolically inert (4).

We next assessed whether the presence of NSCLC altered local metabolite levels. A direct comparison of unadjusted metabolite concentrations in veins draining healthy and tumor-containing lobes showed substantial overlap, with no significant differences (**Supplemental Figure 2B-C**). However, normalizing by each participant’s pulmonary artery values revealed consistent trends in metabolite level changes distinguishing cancerous from healthy lobes (**Figure 1F-H, Supplemental Figure 2D**). These patterns persisted when restricting analysis to participants who had not received neoadjuvant therapy (n = 15) (**Supplemental Figure 3A-C**). These findings highlight both the sensitivity of this intraoperative sampling approach and the challenges in identifying metabolic biomarkers when using a distant and diluted sampling site (for example, peripheral veins) given the small differences in concentrations appreciated in the pulmonary venous system.

Lactate emerged as one of the top metabolites secreted at higher levels in tumor-bearing lungs compared to healthy lungs, while glucose consumption was similar between the groups (**Figure 1I**). The tricarboxylic acid cycle (TCA) cycle metabolites succinate and fumarate were elevated or trended higher in the tumor-draining vein, and glycine was also significantly secreted by tumor-containing tissue. Aspartate and asparagine both showed a trend toward increased secretion by cancerous tissue. These differences were less pronounced when analysis was limited to participants who had not received neoadjuvant therapy, although increased lactate secretion by tumor-bearing lung remained significant (**Supplemental Figure 3D**). Analysis of lipid and metal species revealed no consistent net differences between tumor and healthy lobe circulation (**Supplemental Figure 3E-F**). Notably, lactate secretion directly correlated with tumor PET avidity, but not with tumor size (**Figure 1J, Supplemental Figure 3G**).

These findings both confirm and also question existing paradigms related to lung cancer metabolism. Recent work has suggested that lactate acts as a key carbon source for NSCLC tumors, taken up by tumors for use in the TCA (5–7). Thus, while lactate carbon could contribute to labeling of TCA cycle metabolites, these data demonstrate that lactate is still net produced by tumor-containing lung tissue, at least in the patients we analyzed. This discrepancy may stem from methodological differences, as isotope-labeling does not directly assess the consumption and secretion of metabolites. Furthermore, accumulation of succinate and fumarate in the vein draining the tumor-bearing lung may reflect reduced TCA cycle activity, consistent with lung tumors shifting towards increased aerobic glycolysis (8).

Although previous studies have highlighted aspartate as a limiting metabolite in cancer cells, particularly under hypoxia or in response to impaired mitochondrial respirations (9), these data suggest that NSCLC tumors may secrete aspartate into the venous circulation. This finding raises the possibility that aspartate availability and utilization are context-dependent.

Glycine is a non-essential amino acid but its synthesis is upregulated in cancer. A pan-cancer analysis of over 10,000 patient samples revealed copy number alterations in genes involved in serine and glycine synthesis in approximately 6% of tumors, including NSCLC, with most alterations representing copy number gains (10). These findings suggest that a subset of NSCLCs exhibit increased glycine synthesis. Although we did not perform genomic profiling of the patient tumors in this study, the observed glycine secretion from tumor-bearing lungs supports the possibility that glycine synthesis is upregulated *in situ*. Further work will be needed to determine whether this reflects underlying genomic alterations or another adaptation.

Our work highlights the value of orthogonal and complementary approaches to characterize cancer metabolism *in situ*. Our insights demonstrate the potential of using intraoperative blood sampling with metabolite measurements to provide information regarding nutrient fluxes across tumor-bearing and other diseased organs.

## Methods

### Patient selection

Patients with resectable biopsy-proven non-small cell lung cancer with tumors greater than 1cm on pre-operative imaging were approached for participation in the general thoracic surgery clinic at University of Chicago Medical Center from July 2021 – May 2023. Patients were excluded if they had a history of inherited metabolic disorders, were currently taking oral steroid medications, or were planned to undergo a sub-lobar resection. Informed consent was obtained in clinic during a pre-operative appointment (IRB: UCMC 20-1696). Patient demographics including age, sex, race/ethnicity, comorbidities, frailty status, and pathologic and radiologic evaluation of tumors were recorded **(Figure 1A; Supplemental Table 1)**.

### Intraoperative sampling

At time of oncologic resection, after the dissection of the hilar vessels, blood samples were obtained by the thoracic surgery team from the pulmonary vasculature and by the anesthesiology team from the systemic circulation. Using a 25-gauge needle, arterial inflow to the lung (the pulmonary artery) and venous outflow from the cancer containing lobe’s pulmonary vein and an adjacent healthy lobe’s pulmonary vein were sampled **(Supplemental Figure 1A-B)**. Simultaneously, the anesthesiology team obtained blood samples from a radial arterial line and a peripheral vein to provide an approximation of the concentration gradient change of metabolites across the systemic circulation.

### Validation of sample processing

A single healthy volunteer provided plasma samples for evaluation of whether plasma metabolite composition is altered by time spent on ice prior to sample processing. A 20-gauge butterfly needle was used to collect 5 mL of blood from the antecubital vein of the volunteer. Four 1 mL aliquots were placed into heparin coated cryovials. Samples were left on ice for 4, 15, 25 and 36 min prior to being centrifuged at 850 x g for 10 min at 4 °C. Plasma was isolated and partitioned into 50 μL aliquots in uncoated Eppendorf tubes prior to flash freezing in liquid nitrogen and stored at -80 °C prior to analysis by liquid chromatography/mass spectrometry (LC/MS).

### Initial processing of intraoperative samples

In the operating room, blood samples were partitioned into 1 mL aliquots in EDTA coated cryovials before being centrifuged at 800 x g for 10 min at 4 °C. Plasma was partitioned into aliquots of 100 μL into 1.5 mL uncoated Eppendorf tubes and flash frozen with dry ice prior to transport and storage at -80 °C prior to analysis by LC/MS.

### LC/MS analysis of polar metabolites

Metabolite quantification in human fluid samples was performed as described previously (3). In brief, 5 μL of sample or external chemical standard pool (ranging from ∼5 mM to ∼1 μM) was mixed with 45 μL of acetonitrile:methanol:formic acid (75:25:0.1) extraction mix including isotopically labeled internal standards. All solvents used in the extraction mix were HPLC grade. Samples were vortexed for 15 min at 4 °C and insoluble material was sedimented by centrifugation at 16,000 x g for 10 min at 4 °C. 20 μL of the soluble polar metabolite extract was taken for analysis of polar metabolites by LC/MS.

Metabolite profiling by LC/MS was conducted on a QExactive benchtop orbitrap mass spectrometer equipped with an Ion Max source and a HESI II probe, which was coupled to a Dionex UltiMate 3000 HPLC system (Thermo Fisher Scientific). External mass calibration was performed using the standard calibration mixture every 7 days. An additional custom mass calibration was performed weekly alongside standard mass calibrations to calibrate the lower end of the spectrum (m/z 70-1050 positive mode and m/z 60-900 negative mode) using the standard calibration mixtures spiked with glycine (positive mode) and aspartate (negative mode). 2 μL of each sample was injected onto a SeQuant® ZIC®-pHILIC 150 x 2.1 mm analytical column equipped with a 2.1 x 20 mm guard column (both 5 mm particle size; EMD Millipore). Buffer A was 20 mM ammonium carbonate, 0.1% ammonium hydroxide; Buffer B was acetonitrile. The column oven and autosampler tray were held at 25 °C and 4 °C, respectively. The chromatographic gradient was run at a flow rate of 0.150 mL min-1 as follows: 0-20 min: linear gradient from 80-20% B; 20-20.5 min: linear gradient form 20-80% B; 20.5-28 min: hold at 80%

B. The mass spectrometer was operated in full-scan, polarity-switching mode, with the spray voltage set to 3.0 kV, the heated capillary held at 275 °C, and the HESI probe held at 350 °C. The sheath gas flow was set to 40 units, the auxiliary gas flow was set to 15 units, and the sweep gas flow was set to 1 unit. MS data acquisition was performed in a range of m/z = 70–1000, with the resolution set at 70,000, the AGC target at 1×10^6^, and the maximum injection time at 20 msec.

Following LC/MS analysis, metabolite identification was performed with XCalibur 2.2 software (Thermo Fisher Scientific) using a 5 ppm mass accuracy and a 0.5 min retention time window. For metabolite identification, external standard pools were used for assignment of metabolites to peaks at given m/z and retention time, and absolute metabolite concentrations determined as previously described (3). To account for individual patient differences, metabolite concentrations in the pulmonary veins (cancer-bearing lobe draining pulmonary vein: PVC; non-cancer-bearing lobe (healthy)-draining pulmonary vein: PVH) were normalized to the patient-specific pulmonary artery (PA) concentration. This was achieved by calculating the difference between each vein’s concentration and the corresponding pulmonary artery concentration before comparing the relative consumption and production of individual metabolites.

### LC/MS lipidomics

Positive ion mode analyses of polar and nonpolar lipids were conducted using an LC/MS system composed of a Shimadzu Nexera X2 U-HPLC (Shimadzu) coupled to an Exactive Plus orbitrap mass spectrometer (ThermoFisher Scientific). 10 μL of human fluid sample was precipitated with 190 μL of isopropanol containing 1,2-didodecanoyl-sn-glycero-3-phosphocholine (Avanti Polar Lipids) as an internal standard. After centrifugation, 2 μL of supernatant was injected directly onto a 100 × 2.1 mm, 1.7-μm ACQUITY BEH C8 column (Waters). The column was eluted isocratically with 80% mobile phase A (95:5:0.1 v/v/v 10 mM ammonium acetate/methanol/formic acid) for 1 min followed by a linear gradient to 80% mobile phase B (99.9:0.1 v/v methanol/ formic acid) over 2 min, a linear gradient to 100% mobile phase B over 7 min, then 3 min at 100% mobile phase B. Mass spectrometry analyses were performed using electrospray ionization in the positive ion mode using full scan analysis over 220 to 1,100 m/z at 70,000 resolution and 3 Hz data acquisition rate. Other mass spectrometry settings were as follows: sheath gas 50, in source collision-induced dissociation 5 eV, sweep gas 5, spray voltage 3 kV, capillary temperature 300°C, S-lens RF 60, heater temperature 300°C, microscans 1, automatic gain control target 1×10^6^, and maximum ion time 100 ms. Lipid identities were determined on the basis of comparison to reference standards and reference plasma extracts and were denoted by the total number of carbons in the lipid acyl chain(s) and total number of double bonds in the lipid acyl chain(s). Ion counts normalized to an internal standard were reported.

### Inductively coupled plasma mass spectrometry (ICP-MS) for metal analysis

Samples were digested with 2 mL/g total wet weight nitric acid (Trace metal grade; Fisher) for 24 h at room temperature (RT), followed by treatment with 1 mL/g total wet weight hydrogen peroxide (Trace metal grade; Fisher) for another 24 h at RT. Samples were diluted with ultrapure water (VWR Chemicals ARISTAR ULTRA) followed by ICP-MS (Perkin Elmer Nexion 2000) using 50 ppb Bismuth as internal standard.

### Statistics

Initial descriptive statistical analysis including comparison of individual metabolite concentrations, and linear regression comparing metabolite concentration to patient characteristics including tumor size and PET avidity was performed in R version 4.2.0 (Foundation for Statistical Computing, Vienna, Austria). Metaboanalyst 6.0 (11) was used to perform pairwise comparisons of metabolite concentrations, partial least squares determinant analysis, create metabolite heat maps, and identify variables of importance.

### Study approval

Written informed consent was obtained from all patients. This study was initially approved on 5/10/2021 and was overseen for the duration of the study protocol by the University of Chicago IRB (IRB#20-1696). All Intraoperative images were obtained with patient consent at time of their procedure and consent has been retained.

### Data availability

Values for all data points in graphs are reported in the Supporting Data Values file. Any additional information required to reanalyze the data reported in this paper is available from the lead contact upon reasonable request.

## Supporting information

Supporting Data Values

## Author contributions

Conceptualization: J.R.K., K.L.A., M.G.V.H., M.L.L.M., P.P.H.; Methodology: J.R.K., K.L.A., M.G.V.H., M.L.L.M., P.P.H.; Investigation: J.R.K., K.L.A., R.N., A.D., N.K.D., M.W., T.K., A.S.Z., N.C., A.S., D.S.B., M.K.F., J.S.D.; Resources: M.G.V.H., M.L.L.M., P.P.H.; Writing – Original Draft: J.R.K., K.L.A., M.G.V.H., M.L.L.M., P.P.H.; Writing – Review & Editing: All authors; Supervision: A.M., Y.M.S., C.B.C., M.G.V.H., M.L.L.M., P.P.H.; Funding Acquisition: M.G.V.H., M.L.L.M., P.P.H.

## Acknowledgements

K.L.A. was supported by the National Science Foundation (DGE-1122374) and National Institutes of Health (NIH) (F31CA271787, T32GM007287). Y.M.S. acknowledges support from the NIH (R01CA148828, R01CA245546). M.G.V.H. acknowledges support from the MIT Center for Precision Cancer Medicine, the Ludwig Center at MIT, and the NIH (R35CA242379, P30CA1405141). Metabolomics work performed in NSCLC patients was supported by the Lung Cancer Research Foundation.

## Figure Legend

**Supplemental Table 1.** Clinical and pathological characteristics of study participants. Summary of demographic, clinical and pathological data of the 20 non-small cell lung cancer resections from 19 patients.

| Participant | Age | Sex | Race | Tumor Location | Histology | Neoadjuvant Therapy | Pathology Tumor Size (cm) | TNM Status Pathologic Stage | Comorbidities |
| --- | --- | --- | --- | --- | --- | --- | --- | --- | --- |
| 1 | 66 | F | Caucasian | RML | Adeno | None | 2.8 | pT1cN0 | Breast Cancer, HLD, HTN |
| 2 | 76 | F | Caucasian | RLL | Adeno | Carboplatin + Pemetrexed | 2.8 | ypT2aN2 | HLD, HTN |
| 3 | 81 | F | Caucasian | RUL | Adeno | None | 3.1 | pT1cN0 | Breast Cancer, Hypothyroidism, Raynaud's disease |
| 4 | 62 | M | Caucasian | RUL | Adeno | None | 3.5 | pT2aN0 | BPH |
| 5 | 67 | M | African American | LLL | Adeno | None | 1.9 | pT1bN1 | Aortic stenosis, CKD, COPD, DM, Gout, HLD, HTN |
| 6 | 70 | F | Caucasian | RML | Adeno | None | 1.8 | pT1bN1 | CAD, Hypothyroid, HLD, HTN, MDD, OA |
| 7 | 74 | F | Caucasian | LUL | Adeno | Clinical Trial: Cisplatin + Pemetrexed +/- Nivolumab | 0 | ypT0N0M0 | DM, HLD, HTN, IBS, TIA |
| 8 | 79 | F | Caucasian | RML | Adeno | None | 0.9 | pT1aN0 | Breast Cancer, Lymphoma, HLD, HTN, hypothyroid |
| 9 | 73 | F | Caucasian | RLL | Adeno | Cisplatin + Pemetrexed | 2.2 | ypT1cN0 | HLD, HTN, OA |
| 10 | 67 | M | Caucasian | LUL | SCC | Clinical Trial: Cisplatin + Pemetrexed +/- Nivolumab | 0.1 | ypT1aN0 | HLD |
| 11 | 69 | M | Caucasian | RUL | Adeno | None | 5.7 | pT3N0 | DM, HTN |
| 12 | 28 | F | African American | LUL | Adeno | None | 1.5 | pT2aN1 | IBS, Iron deficiency anemia, Li-Fraumeni syndrome |
| 13 | 76 | F | Caucasian | LUL | Adeno | None | 2.6 | pT1cN0 | Age Related Macular Degeneration, DM, HTN, HLD |
| 14 | 77 | F | African American | LLL | Adeno | None | 1 | pT3N0 | Diverticulosis, HLD, HT |
| 15 | 73 | F | African American | RLL | SCC | None | 3.5 | pT2aN2 | Anxiety, GERD, HTN, MDD, OA |
| 16 | 77 | M | Caucasian | LLL | Adeno | None | 1.5 | pT1bN0 | HLD, Rectal Adenocarcinoma |
| 17 | 67 | F | Caucasian | LUL | Adeno | Carboplatin + Paclitaxel + Pembrolizumab | 2.9 | ypT3N0 | CAD, GERD, HLD, Thrombocytopenia |
| 18* | 47 | F | Other | LUL | Adeno | None | 3.2 | pT1bN0 | DM, HLD, HTN, OA, OSA |
| 19* | 47 | F | Other | RLL | Adeno | None | 3.8 | pT2aN0 | DM, HLD, HTN, OA, OSA |
| 20 | 49 | F | Caucasian | RUL | Adeno | None | 2.3 | pT1cN0 | Asthma, GERD, HTN |
| Abbreviations: M – Male; F- Female; RUL – Right Upper Lobe; RML – Right Middle Lobe; RLL – Right Lower Lobe; LUL – Left Upper Lobe; LLL – Left Lower Lobe; Adeno – Adenocarcinoma; SCC – Squamous Cell Carcinoma; BPH – Benign Prostatic Hypertrophy; CAD – Coronary Artery Disease; COPD – Chronic Obstructive Pulmonary Disease; CKD – Chronic Kidney Disease; DM – Diabetes Mellitus; GERD – Gastroesophageal Reflux Disease; HLD – Hyperlipidemia; HTN – Hypertension; IBS - Irritable Bowel Syndrome; MDD – Major Depressive Disorder; OA -Osteoarthritis; OSA – Obstructive Sleep Apnea; TIA – Transient Ischemic Attack. |  |  |  |  |  |  |  |  |  |
| * Subject 18/19 was the same patient with co-synchronous bilateral tumors that were resected in separate encounters |  |  |  |  |  |  |  |  |  |

## Supplemental Figure Legends

**Supplemental Figure 1.**
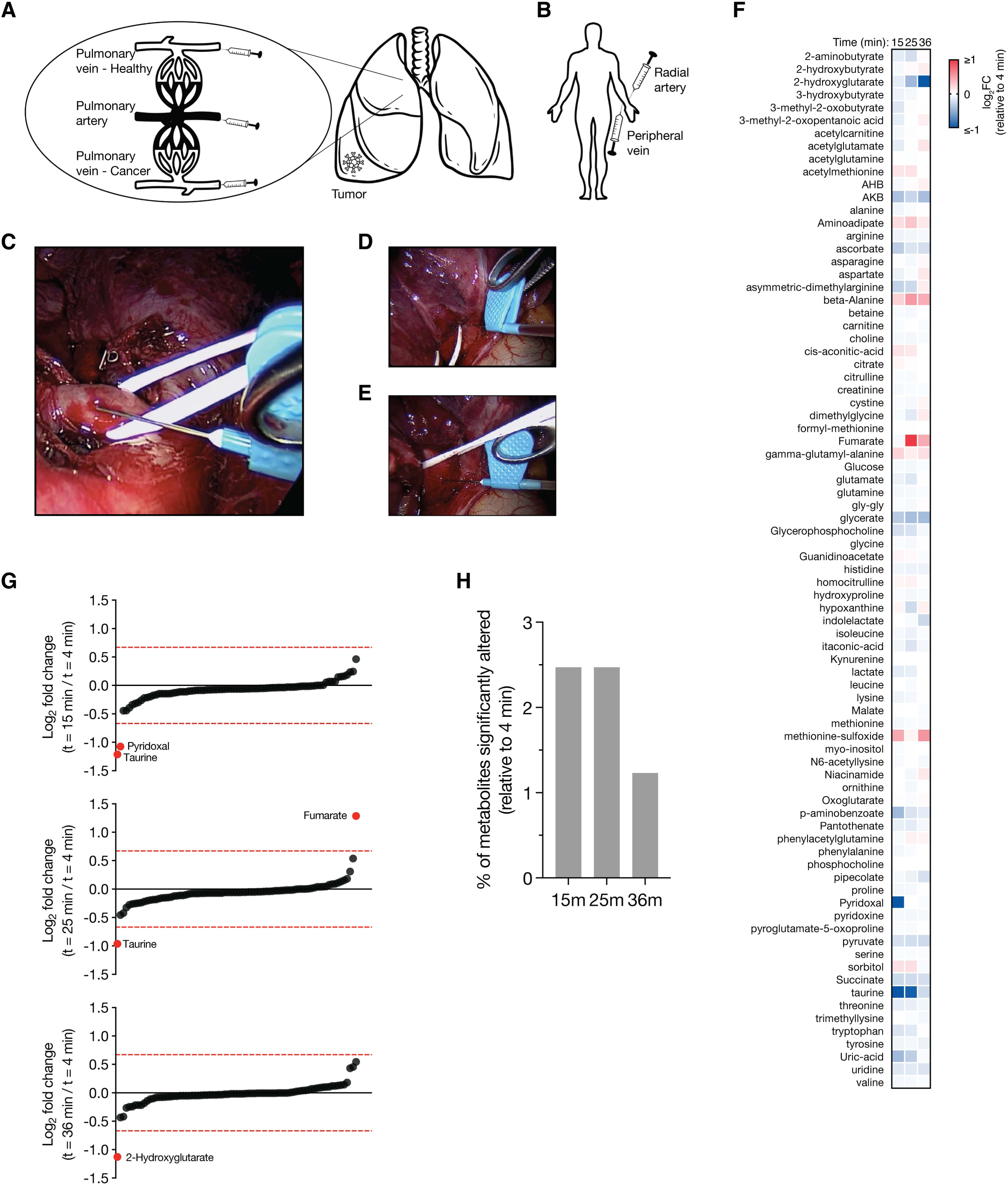
Validation of intraoperative blood sampling and processing conditions. **(A-B)** Schematic illustrating sampling sites in the study. (A), Pulmonary vasculature including tumor-bearing and non-tumor-bearing (healthy) lung lobes. (B), Systemic sites including the radial artery and a peripheral vein. **(C-E)** Intraoperative imaging during a video assisted thoracoscopic right upper lobectomy. Sampling of the pulmonary artery (C), pulmonary vein draining the cancer containing lobe (right superior pulmonary vein) (D); and healthy lobe (right middle pulmonary vein) (E) is shown. **(F)** Heatmap showing average log₂ fold change in plasma metabolite concentrations over the sample processing time measured relative to baseline (t = 4 min). AHB: alpha-hydroxybutyrate; AKB: alpha-ketobutyrate; gly-gly: glycyl-glycine. **(G)** Plot showing ranked metabolites based on absolute log₂ fold change at different times during sample processing as indicated. Metabolites exceeding the significance threshold (|log₂ fold change| > 0.58) at each time point are highlighted in red. **(H)** Bar plot showing the percentage of metabolites significantly altered (|log₂ fold change| > 0.58) at each time point during sample processing relative to t = 4 min.

**Supplemental Figure 2.**
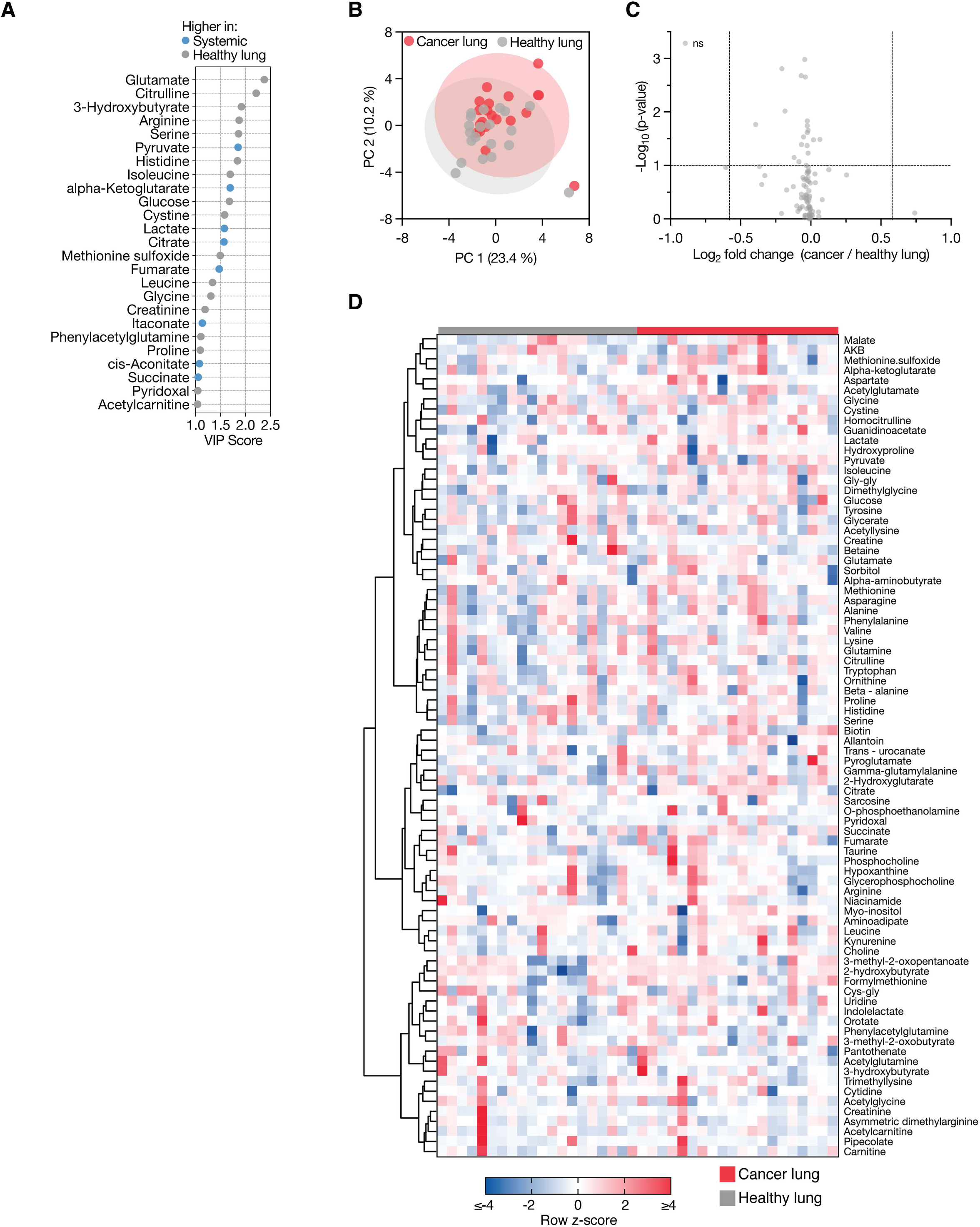
Direct comparison of pulmonary vein metabolite levels lacks sensitivity to distinguish cancer from healthy lung. **(A)** Variable importance in projection (VIP) plot of the top 25 metabolites distinguishing non-tumor-bearing (healthy) lung circulation (pulmonary vein of a non-tumor lobe minus pulmonary artery) from systemic circulation (peripheral vein minus radial artery). Metabolites in blue are decreased in healthy lung relative to systemic circulation while those in grey are elevated. Derived from data shown in Figure 1B. **(B)** Partial least squares discriminant analysis (PLS-DA) comparing metabolite profiles from the pulmonary vein of a healthy lobe versus that of a cancer-bearing lobe (n = 20). **(C)** Volcano plot depicting metabolites significantly altered between cancer-bearing and healthy pulmonary veins. Significance was defined as a fold change >1.5 and a raw p-value <0.1 (paired two-tailed t-test, n = 20). **(D)** Hierarchical clustering of z-score–normalized metabolite concentrations in blood from cancer-bearing versus healthy lung lobes (both calculated as pulmonary vein minus pulmonary artery). Rows represent individual metabolites; patient groups were not clustered (n = 20). AKB: alpha-ketobutyrate; cys-gly: cysteinylglycine; gly-gly: glycyl-glycine.

**Supplemental Figure 3.**
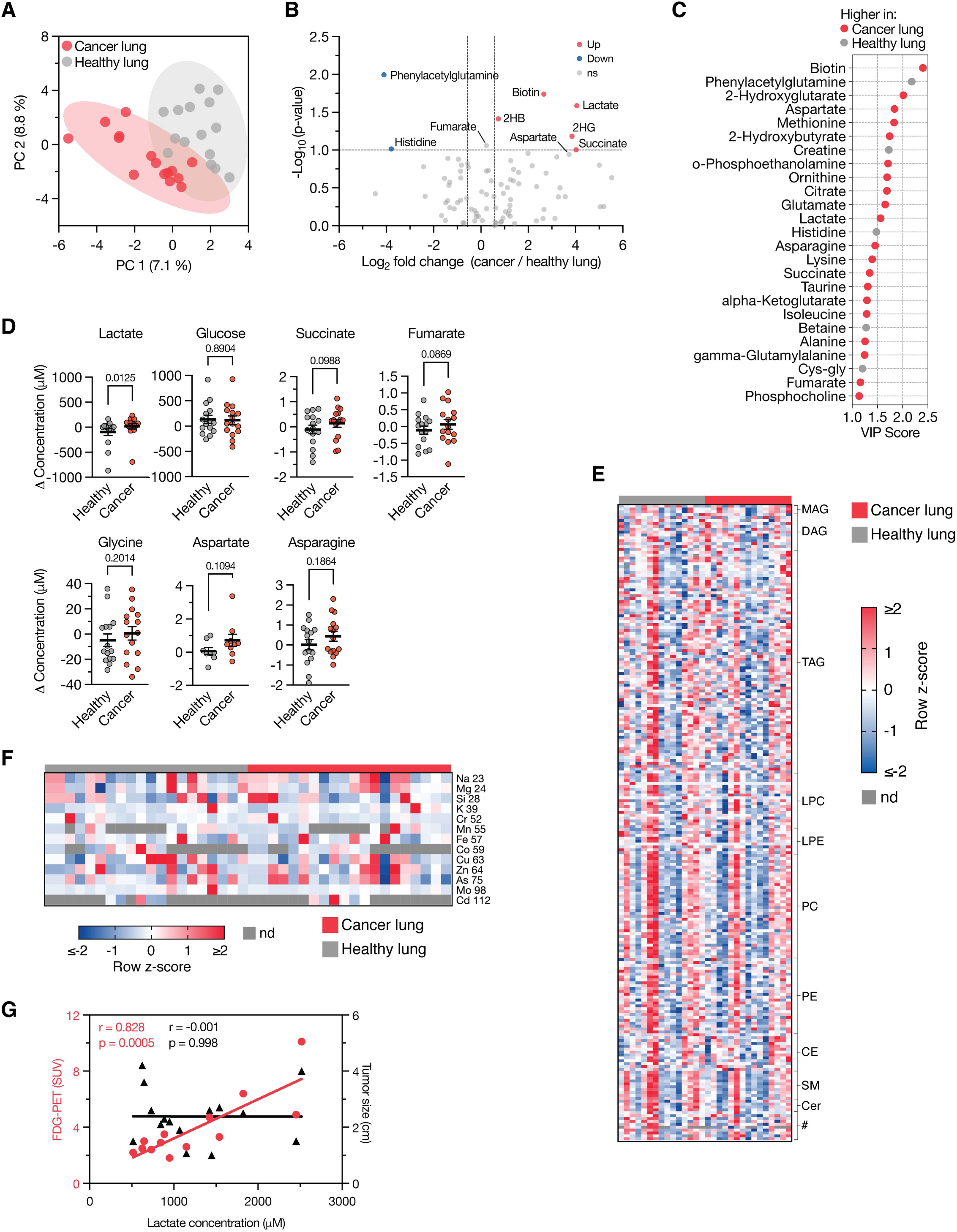
Tumor-associated metabolic alterations are found in patients who did not receive neoadjuvant therapy. **(A)** Partial least squares discriminant analysis (PLS-DA) comparing metabolite profiles from cancer-bearing versus non-cancer bearing (healthy) lung tissue (both calculated as pulmonary vein minus pulmonary artery) in patients who did not receive neoadjuvant therapy (n = 15). **(B)** Volcano plot showing metabolites significantly altered between cancer-bearing and healthy lung in this subset of patients. Significance defined as fold change >1.5 and raw p < 0.1 (paired two-tailed t-test, n = 15). 2HB: 2-Hydroxybutyrate; 2HG: 2-Hydroxyglutarate. **(C)** Variable importance in projection (VIP) plot of the top 25 metabolites distinguishing cancer from healthy lung in patients without neoadjuvant therapy. Metabolites in red are elevated in cancer; those in grey are elevated in healthy lung. Based on data in panel (A). **(D)** Selected metabolites differing between cancer-bearing and healthy lung in patients who did not receive neoadjuvant therapy. Data are mean ± SEM; p-values from Wilcoxon matched-pairs signed rank test (n = 15). **(E)** Heatmap of lipid species in the peripheral vein draining healthy (grey) or cancer-bearing (red) lung, normalized to the pulmonary artery. Each row is z-score normalized across samples (n = 15). Lipid classes: MAG, monoacylglyerol; DAG, diacylglycerol; TAG, triacylglycerol; LPC, lysophosphatidylcholine; LPE, lysophosphatidylethanolamine; PC, phosphatidylcholine; PE, phosphatidylethanolamine; CE, cholesteryl ester; SM, sphingomyelin; Cer, ceramide. @ denotes the following metabolites (top to bottom): sphingosine, palmitoylethanolamide, cholesterol, campesterol, piperine, coenzyme Q9, coenzyme Q10, alpha-tocopherol, delta-tocopherol, gamma-tocopherol. **(F)** Heatmap of metal ion concentrations in the pulmonary vein draining healthy (grey) and cancer-bearing (red) lung lobes, normalized to the pulmonary artery. Data represent all participants (n = 20). Each row is z-score normalized across samples. **(G)** Scatter plot showing lactate levels in the cancer-draining pulmonary vein plotted against FDG-PET SUV (red, n = 13) or tumor size size (black, n = 15) in patients that did not receive neoadjuvant therapy. Pearson correlation coefficients and p-values are indicated.

